# Effects of intra-hippocampal corticosterone and sleep on consolidation of complex memory of aversive experience in rats

**DOI:** 10.1101/2021.08.30.458150

**Authors:** Alena Brukhnová, Ewa Szczurowska, Čestmír Vejmola, Rachel R. Horsley, Eduard Kelemen

## Abstract

Formation and consolidation of memories for highly stressful (traumatic) events is a complex process that involves interplay between multiple memory systems and has implications for etiology and treatment of stress- and trauma-related disorders. Here we study effects of sleep/wake state and high intra-hippocampal corticosterone on consolidation of aversive contextual memories as well as consolidation of association between simple trauma-related cues and fear response in rats. Animals were implanted with EEG and EMG electrodes for sleep assessment and cannulas for intra-hippocampal corticosterone application. They were familiarized to a “safe box” and then trained in fear conditioning paradigm in a distinct “shock box” with a prominent simple auditory cue serving as a phasic background cue. Immediately after conditioning, animals received bilateral intra-hippocampal saline (1μl) or corticosterone (10ng in 1μl saline) injection and were either allowed to sleep or were kept awake for a following two-hour consolidation period. Memory test twenty-four hours later revealed that the saline-injected animals with sleep during consolidation had significantly stronger freezing response in the shock box compared to the safe box as well as increased freezing in response to the tone. Lack of post-learning sleep in saline injected animals led to generalization of fear response to the safe context, while association between simple cue and fear response was preserved. High intra-hippocampal corticosterone level during memory consolidation led to generalization of fear response to the safe context, regardless of sleep/wake state, while enhancement of response to single stimulus was not observed. Our results show how manipulation of conditions during consolidation can lead to greatly variable complex memories for a traumatic episode and distinct behavioral outcomes.

**Highlights:**

- We studied effect of sleep and intrahippocampal corticosterone on consolidation of memories surrounding stressful event modeled by fear conditioning in rats.
- Sleep following traumatic fear conditioning event is important for subsequent manifestation of fear response (freezing) specifically in the context of traumatic event but not in a neutral safe context.
- Lack of sleep or high intra-hippocampal corticosterone level during memory consolidation leads to generalization of fear response to both the traumatic and safe context.
- Increased freezing in response to a trauma-related auditory cue was observed in saline injected rats regardless of wake/sleep state during consolidation.
- Post-learning intra-hippocampal corticosterone injection blocked response to a trauma-related auditory cue regardless of wake/sleep state during consolidation.

## 1. Introduction

Memories created around stressful life events play a key role in etiology of certain mental disorders, such as post-traumatic stress disorder (PTSD) and other trauma-related, anxiety and mood disorders. Among the effects of traumatic experience on memory is impairment of memories for the spatiotemporal context of traumatic episode (Dancu et al., 1996; Ursano et al., 1999; Layton & Krikorian, 2002). This may manifest as overgeneralization of fear responses, i.e. the activation of fear and anxiety in safe situations (Kaouane et al., 2012; Kaczkurkin et al., 2017). Another frequent memory effect of traumatic experience is strong association of fear/anxiety response with simple cues (auditory, visual, olfactory) present around the time of traumatic experience. These cues can than develop into so-called “trigger cues” for recollection (flashbacks) of traumatic event.

Formation of long-term memories strongly depends on conditions surrounding memory consolidation processes, with particularly strong influence of sleep/wake state and stress hormone levels. Sleep benefits of consolidation of hippocampus-dependent (Phillips and LeDoux, 1992; 1994) contextual information has been well established (Oyanedel et al., 2014; Kelemen et al., 2014; Inostroza and Born, 2013; Diekelmann and Born, 2010; Wagner and Born, 2008). High densities of glucocorticoid and mineralocorticoid receptors in the hippocampus (Joëls, 2008) make hippocampus-dependent memories particularly sensitive to increased corticosteroid levels following stressful events (Lupien & Lepage, 2001; Maheu et al., 2005; Rimmele et al., 2013). Some previous studies reported enhancing effects of glucocorticoids (GCs) on memory consolidation (Cottrell and Nakajima, 1977; Micheau et al., 1985; Roozendaal and McGaugh, 1997; deQuervain et al., 2009), however this effect is dependent on wake or sleep conditions of the subject. We and others have previously shown in *emotionally neutral* situations that while corticosterone (in rats) or cortisol (in humans) impairs memory consolidation in sleep, in wakefulness GCs enhance otherwise poor consolidation of hippocampus-dependent memories (Wilhelm et al, 2011; Kelemen et al., 2014). Corticosterone in the hippocampus affects not only consolidation of contextual cues of stressful experience, but also conditioning to co-occurring prominent simple auditory phasic background cue (Kaouane et al., 2012), thus creating a complex pattern of memory alternations involving multiple memory systems. How effects of sleep/wake state and corticosterone in combination affect consolidation of complex memories for *traumatic* experience remains an open question.

Against this backdrop, we used fear conditioning paradigm in rats to study the complex, integrated effects of sleep and increased intra-hippocampal corticosterone levels on consolidation of memories for a *context* of traumatic (fear conditioning) event and memories for *auditory background cue* present during traumatic experience in rats.

## 2. Materials and methods

Forty male adult Long-Evans rats 3-5 months old at the time of the experiment were used. The rats were housed in standard plastic boxes with free access to food and water in controlled 12 hours light/12 hours dark cycle with lights on at 6 AM. The room temperature was maintained at 20-24°C and humidity between 30-70%. All rats were experimentally naive and tested once. All animal procedures were approved by the committee for the ethical treatment of animals and animal welfare at the National Institute of Mental Health (Project No. 13/2016) and complied with the Animal Protection Act of the Czech Republic and European Union directive 2010/63/EC.

Electrodes for electroencephalography (EEG) recording were made of bone screws attached to Mill-Max connector through insulated nickel-chromium wires (75 μm in diameter, California Fine Wire Company, Grover Beach, CA, USA). A wire with free end stripped of insulation served as an electromyography (EMG) electrode. Intra-hippocampal cannulas for corticosterone injections were prepared of 22-gauge stainless steel tube following procedures described in Klement et al. (2005). Prior to electrode and cannula implantation, the rats were anesthetized by isoflurane and placed in the stereotaxic frame (Stoelting, Wood Dale, IL, USA), during the surgery they were kept under general anesthesia (2.5% isoflurane). After the scalp was removed and the skull was exposed, five holes were drilled for the electrode screws. Three screws, later used for EEG recording, were inserted in the frontal and parietal areas of the skull, the other two screws, in the occipital bone, were used as grounding and reference electrodes. The EMG electrode was inserted under the skin into the dorsal neck region. Two holes for the cannulas were drilled bilaterally (3 mm posterior, 2.5 mm lateral from Bregma). Two cannulas were inserted 2 mm below the surface of the skull just above the dorsal hippocampus surface. The cannulas and the Mill-Max connector were fixed to the skull with dental cement (Dentalon, Kulzer GmbH, Hanau, Germany). Removable wires were placed inside the guide cannulas to prevent infection and to avoid clogging. During experiment the wires were removed for corticosterone or saline injection. The wound was sutured and treated with povidone-iodine (Betadine 2%, EGIS, Prague, Czechia) and antibiotics ointment to prevent inflammation (Framykoin, Zentiva Group, Prague, Czechia). Before beginning of the experiment, the animals were allowed to recover for at least one week until the wound has fully healed.

After recovery from the surgery, the experiment resumed (Fig. 1A). Two boxes (45 × 45 cm) were used as a “safe box” and “shock box” respectively. They were made of different materials (plexiglass vs. OSB board) and were painted differently (light and dark vertical stripes vs. all grey walls). The identity of “shock” and “safe” box was alternated between rats. The “safe box” had a plywood floor, the “shock box” had a conductive grid that was connected to the Arduino-controlled electric shock delivery system.

**Figure 1.**
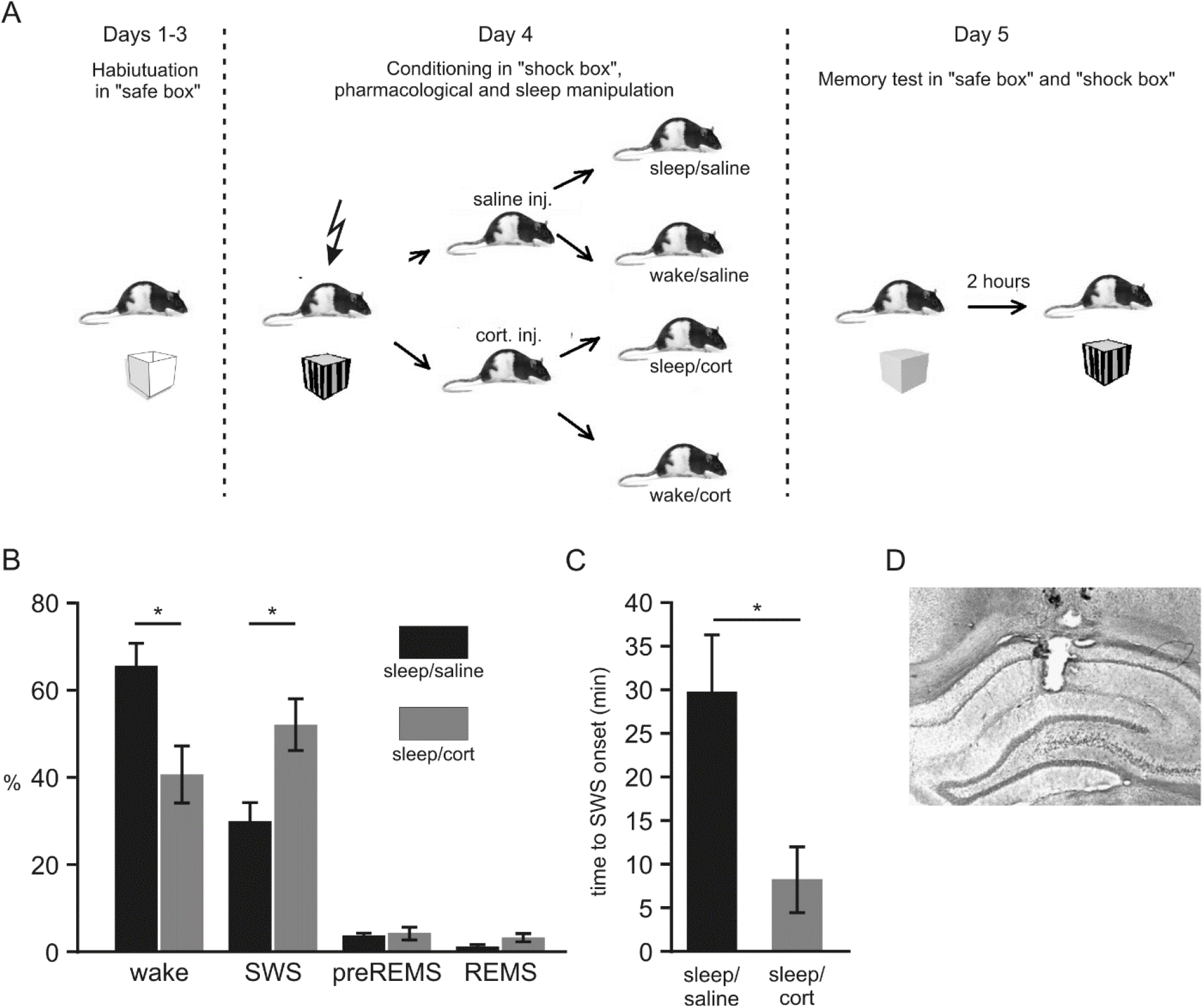
A) Experimental protocol. Days 1-3: Habituation in a *safe box* and a *sleep box*. Day 4: Contextual conditioning in a *shock box* followed by saline or corticosterone intra-hippocampal injection. During the following two-hour period in the sleep box the animals were either kept awake, or were allowed to sleep. Day 5: Memory test in the safe box and shock box. B) Percentage of time spent in different sleep stages during two-hour consolidation period in sleep/saline and sleep/cort groups. C) SWS onset during the consolidation period for the two sleeping groups. D) An example of the histological preparation showing position of a cannula in the dorsal hippocampus. (Values are shown as mean ± SEM; * p<0.05.)

All experimental manipulations were performed between 10:00 AM and 4:00 PM. For three days animals were daily habituated to the “safe box” for four minutes and to the third, distinct “sleeping box” in a different room for at least two hours. The fear conditioning training took place on the 4th day of the experiment. The animals spent 5 minutes and 20 seconds in the “shock box”. During this period the animals received two electric shocks (0.5 mA for 1 s) to their feet and twice a tone (15 s, 65 dB, 1 kHz) was presented. The conditioning paradigm had the following sequence: pause 60 s – foot shock 1s – pause 80 s – tone 15 s – pause 30 s – tone 15 s – pause 90 s – foot shock 1s – pause 30 s. Note that the tones did not directly precede and predict the shocks (Kaouane et al., 2012). Immediately after the conditioning, 30-gauge injection cannula was inserted into a guide cannula 3.4 mm deep into the brain and the animals received bilateral injection of 10ng of corticosterone (corticosterone-HBC (2-hydroxypropyl-b-cyclodextrin) complex, Sigma-Aldrich, St. Louis, MO, USA) in 1µl of saline or equivalent volume of saline solution into dorsal hippocampi. Then they were transferred to the sleep box where they were recorded for two hours or they were kept awake using gentle handling procedure (Kelemen et al., 2014). This manipulation lead to four experimental groups with different consolidation conditions: sleep/saline, sleep/cort, wake/saline and wake/cort.

Twenty-four hours after conditioning, the behavior was tested. First, the rats spent two minutes in the “safe box”. After one minute of exploration the tone was played (with the same parameters as during the training). Two hours later (which animals spent in their home cages), the animals were exposed to the “shock box” for two minutes. The animal’s position was recorded and stored for off-line analysis. Freezing time in the two boxes was measured to assess the fear response. Right after the test, animals were sacrificed (deeply anesthetized with ether and decapitated). Their brains were collected and processed for histological evaluation of position of the cannulas (Fig. 1D).

During the two-hour post training consolidation period, the rats were connected to recording system (Axona Ltd. St. Albans, U.K.). EEG signals were filtered between 0.1 and 300 Hz and recorded at 2000 samples per second. EMG signals were filtered between 30 and 300 Hz and sampled at 2000 Hz. Recordings were evaluated off-line using a custom made Matlab (MathWorks, Natick, MA, USA) program (E. Kelemen). Each 10 second interval of EEG and EMG recordings was manually scored. According to the EEG and EMG signals, the sections were divided into individual phases – slow-wave sleep (SWS), rapid eye movement sleep (REMS), preREM sleep, or awake. The total time spent in each phase and the time to sleep onset was calculated.

The evaluation of animals’ behavior was performed using Ethovision XT 11.5 (Noldus, Wageningen, Netherlands) and custom made Matlab code (E. Kelemen). Freezing was detected if the position of an animal did not change for more than two seconds (Fig. 2A). To assess the contextual fear response we compared freezing during the first 60 seconds in the safe box with the first 60 seconds in the shock box. To assess response to the tone, freezing during the 15 seconds of tone was compared to 15 seconds preceding the tone onset in the safe box.

**Figure 2.**
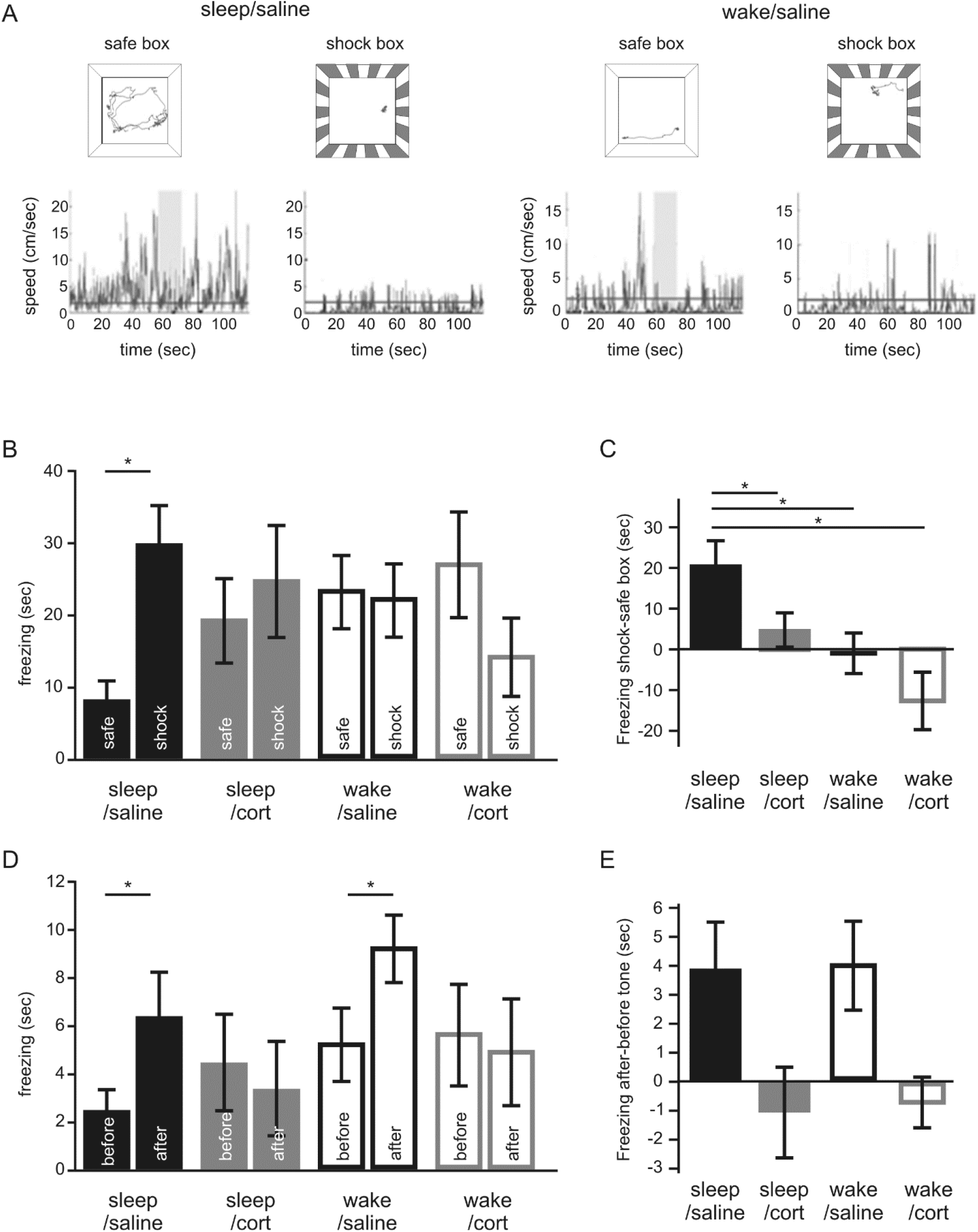
Effects of sleep and intra-hippocampal corticosterone on contextual conditioning. A) Behavior in safe box and shock box during test for an example animal from sleep/saline group (left) and an animal from wake/saline group (right). The upper part of the panel shows rat’s trajectory during test in safe box and shock box. The lower part shows the calculated rat’s speed of movement in two different environments. Horizontal line at 2 cm/sec is the threshold for freezing. Grey rectangle marks the time when the tone was played. B) Freezing time in the safe box and shock box during the first 60 s in both environments. C) Difference of freezing time in shock box and safe box. D) Freezing time in the safe box during fifteen-second intervals before and after the tone. E) Difference in freezing time after and before the tone. (Values are shown as mean ± SEM; * p<0.05.)

For statistical assessment SPSS Statistics package (IBM, Armonk, NY, USA) was used. We used paired t-tests and two-way ANOVAs with difference in freezing as dependent variables and sleep/wake conditions and corticosterone/saline injection as independent variables.

## 3. Results

As a prerequisite for main analysis, we first assessed the effect of intra-hippocampal corticosterone injection on sleep during the two-hour post-conditioning consolidation period. Animals with intra-hippocampal corticosterone injection (sleep/cort) spent less time awake than animals after saline injection (sleep/saline) (t(12) = 2.747, p = 0.018; Fig. 1B). This was the result of increased time in SWS after corticosterone (t(12) = 2.777, p = 0.017). The time spent in REMS and pre-REM sleep was not significantly different between sleep/cort and sleep/saline groups (p’s > 0.05). Further analysis indicated that the increased time in slow wave sleep after corticosterone was caused by increased length of slow wave sleep intervals, rather than their increased number. While the length of SWS intervals was significantly longer in sleep/cort group compared to sleep/saline group (t(12) = 2.28, p = 0.04), the number of SWS intervals was not different (t(12) = 0.85, p = 0.41). Sleep onset was assessed as time to first at least 30-second-long continuous period of SWS. Animals of sleep/saline group took longer to fall asleep compared to sleep/cort group (t(11) = 2.766, p = 0.018; Fig. 1C). All these analyses point to higher sleep pressure after intra-hippocampal corticosterone injection. These analysis also showed that both groups spent significant part of post-conditioning period asleep, which puts us in position to assess effects of sleep and intra-hippocampal corticosterone on contextual fear conditioning.

To analyze spatial context-driven fear responses, we assessed freezing during the first 60 seconds in the shock box and safe box during memory test (day 5 of experiment) in all four experimental groups (Fig 2B). The difference in the freezing in shock box and safe box was calculated for each consolidation condition and compared (Fig. 2C). Two-way ANOVA indicated significant effect of sleep (F(1,36) = 9.939, p = 0.003), significant effect of corticosterone (F(1,36) = 4.587, p = 0.039) and no interaction (F(1,36) = 0.120, p = 0.731). Post hoc tests showed that sleep/saline group differed significantly from each of the other conditions (p’s < 0.05). In sleep/saline group, the total freezing time in the shock box was significantly longer than the freezing time in the safe box (t(9) = 3.126, p=0.012). For the other three consolidation conditions there was no significant difference between freezing times in the two boxes (Fig. 2B). Altogether, these analyses show context-specific fear reaction in the shock box, but not safe box, in the sleep/saline group. In the other three consolidation conditions, generalization of fear response to the safe spatial context was observed.

We next analyzed the rat’s fear responses to the phasic background auditory cue present during conditioning. During test in the safe box, the freezing during 15 seconds before the tone and 15 seconds while the tone was on was analyzed (Fig. 2D). The difference in freezing after and before the tone onset was calculated for each condition and compared (Fig. 2E). Two-way ANOVA indicated significant effect of corticosterone (F(1,33)=8.832, p=0.005), no significant effect of sleep (F(1,33)=0.019, p=0.890) and nonsignificant interaction (F(1,33)=0.007, p=0.934). In sleep/saline group, the total freezing time before the tone was significantly shorter than the freezing after (t(9)=2.281, p=0.048). Similarly in wake/saline group, the total freezing time before the tone was significantly shorter than the freezing after (t(11)=2.458, p=0.032). For the other two consolidation conditions there was no significant difference between freezing before and after the tone (Fig. 2D). These results suggest that the tone enhanced the fear response in saline injected rats, whether they slept or remained awake during post-training consolidation period. In contrast, corticosterone injected rats did not increase freezing after the tone, regardless of their sleep/wake conditions.

## 4. Discussion

We studied effects of two important factors (sleep/wake state and intra-hippocampal corticosterone) on consolidation of traumatic experience in rats. Since overall memory of an event and its effect on future behavior of an animal results from combination of different partial memories mediated by functionally and anatomically distinct memory systems, it is beneficial to study relevant memory systems in combination. We focused on interaction between contextual conditioning and phasic background cue conditioning as two key components of integrated memory for traumatic event. In sleep/saline conditions, good contextual discrimination combined with reliable association between sound and fear response. These conditions appeared optimal for creating full representation of the event. In wake/saline conditions fear response generalized to the safe context, while sound triggered even more enhanced fear response. Of the four consolidations conditions studied, this one is most likely to lead to de-contextualization and exaggerated manifestation of fear memory, and increased likelihood of “inappropriate” or even pathological activation of fear response in safe environments. Intra-hippocampal corticosterone (in sleep or wake) was associated with generalization of fear response to safe context, without additional fear response evoked by the sound. Although contextual memory appeared impaired in these conditions, lack of increased fear reaction to sound may compensate for contextual overgeneralization and ameliorate over activation of fear response.

Our findings are consistent with the notion that sleep benefits consolidation of hippocampus-depended memories (Oyanedel et al., 2014; Kelemen et al., 2014; Inostroza and Born, 2013; Diekelmann and Born, 2010; Wagner and Born, 2008). The effects of glucocorticoids in combination with sleep/wake conditions on memory consolidation have been studied earlier in the context of emotionally neutral learning paradigms. While in emotionally neutral situations, increased corticosterone levels helped consolidation of hippocampus-dependent spatial information in awake animals (Kelemen et al., 2014) and humans (Wilhelm et al, 2011); the current study showed that in a stressful (traumatic) situation in rats, consolidation of contextual information was impaired. The discrepancy between these observations can be a consequence of 1) different emotional valence and therefore different stress levels in the different experimental paradigms. Unlike in low-stress learning, in stressful conditions other brain regions, especially the amygdala, are affected and interact with the hippocampus (de Quervain et al., 2009), which may impair consolidation in stressful conditions. On the other hand, absence of amygdala input in stress-free learning may result in enhancement of contextual memory consolidation by intra-hippocampal corticosterone in wakefulness (as seen in Kelemen et al., 2014 and Wilhelm et al, 2011). 2) Another potential source of curious inconsistency between Kelemen et al. (2014) and the current study could be different learning paradigms. In the previous study, we studied memory for object location; in the current study we tested memory for context-fear association.

There are at least two (not mutually exclusive) ways to think about the overgeneralization of fear that we observed. According to one hypothesis, *error in contextual discrimination*, i.e., inability to differentiate the safe context from the dangerous context, is the primary cause for overgeneralization. Facts that 1) the hippocampus is encoding spatial context, 2) this structure is endowed with pattern separation functions, and 3) our manipulation targeted the hippocampus all make the error in contextual discrimination hypothesis plausible. We have previously shown that stressful experience reactivates consolidated memories even outside the context of their acquisition and these reactivations are hippocampus-dependent (Jezek et al., 2010). In the context of our current experiment, this raises the possibility of reactivation of one-day-old memories of the safe context due to increased corticosterone in the hippocampus. This reactivated memory could then improperly associate with the fear response leading to manifestation of fear in the safe context. According to an alternative hypothesis, *error in emotional tagging* may be responsible for overgeneralization. In this view, discrimination of the different contexts is intact, but inadequately contextualized fear response is expressed in different contexts. Further studies of hippocampal neuronal dynamics under stress and high corticosterone concentration will be needed to test and disentangle the two hypotheses.

Trauma-related auditory cue triggered increased freezing in saline injected animals and not in corticosterone injected animals, regardless of sleep/wake state. Absence of sleep effect is consistent with the notion that hippocampus-independent simple cue conditioning memory is not dependent on sleep (Graves et al., 2003). While in previous studies tone directly preceded the shock during conditioning (Graves et al., 2003) in our experiment the tone was not directly associated with the shock and served rather as a phasic cue in a “background conditioning” paradigm (Majchrzak et al., 2006; Kaouane et al., 2012). The detrimental effect of intra-hippocampal corticosterone on hippocampus-independent tone-fear association is paradoxical, and must be interpreted carefully, since in corticosterone groups the freezing in the safe box was increased due to contextual overgeneralization, which could have hindered detection of the tone effect. Yet, we may speculate that our observations are result of a negative feedback loop triggered by activation of hippocampal corticoid receptors (de Kloet et al., 1998) that decreased corticosteroid milieu in other brain regions and affected consolidation of memory for association between the tone and fear response. Similar negative feedback effect may explain observed increased sleep in response to corticosterone (Fig. 1B, 1C), which is consistent with reports of increased time of SWS after social defeat stress in mice (Fujii et al., 2019).

Of the two categories of corticosterone receptors in the hippocampus, mineralocorticoid receptors with higher affinity to corticosterone are prone to activation even in basal conditions and glucocorticoid receptors with lower affinity activate predominantly under increased stress (de Kloet et al., 1998; Joëls, 2008). In our experimental paradigm the effect of corticosterone injection was most likely mediated predominantly through glucocorticoid receptors. How physiological effects of corticosterone in the hippocampus translate to impairments of contextual memory consolidation remains an open question. Corticosterone-induced decrease in the incidence of hippocampal sharp waves (Weis et al., 2008), which are critical for memory consolidation (Girardeau et al., 2009) may be one factor behind the effect on memory consolidation.

Impaired memories for spatiotemporal context of traumatic events are among the symptoms of PTSD (Dancu et al., 1996; Layton and Krikorian, 2002). Memory-related symptoms, such as overgeneralization of anxiety to safe contexts or situations, are also present in other anxiety and mood disorders. This makes study of mechanisms of consolidation of context of traumatic memories important from clinical perspective. From a practical point of view, conditions during consolidation may be available for manipulation after the otherwise often unexpected and unpredictable traumatic event. Our experiments show that conditions during consolidation play an important role in determining the fate of traumatic memories and post-event manipulations can thus in principle be effective in ameliorating pathological memory impairments following traumatic experience. Detailed knowledge of multi-factorial effects of sleep/wake state and stress hormones on consolidation of traumatic experiences is crucial for better understanding of etiology of stress-related and anxiety disorders and will lead to much needed novel prophylactic and therapeutic options.

## Authorship contribution statement

Alena Brukhnová: Investigation, Formal analysis, Visualization, Methodology, Writing - original draft, Writing - review & editing. Ewa Szczurowska: Data curation, Formal analysis, Writing - review & editing. Čestmír Vejmola: Methodology. Rachel Horsley: Methodology, Writing - review & editing. Eduard Kelemen: Conceptualization, Software, Methodology, Writing - original draft, Writing - review & editing, Supervision, Funding Acquisition.

## Acknowledgements

The authors wish to acknowledge Iveta Vojtěchová and Dr. Tomáš Petrásek and Dr. Alexandre Seillier for useful comments on the manuscript.

